# Improved long-transcript representation in Oxford Nanopore direct RNA sequencing with UltraMarathonRT

**DOI:** 10.64898/2025.12.10.693495

**Authors:** George Maio, Li-Tao Guo, Sara Olson, Brenton R. Graveley, Jason G. Underwood

## Abstract

While most RNA-seq methods sequence amplified cDNA molecules, the advent of direct RNA sequencing (DRS) empowered the scientific community to read native RNA. This technology unlocked characterization of natural RNA modifications and long RNA isoforms without the inherent biases of PCR amplification. In the library preparation prior to Oxford Nanopore (ONT) sequencing, polyadenylated RNAs are copied by a reverse transcriptase (RT) to generate an RNA-cDNA hybrid. The step aims to eliminate the secondary and tertiary structure inherent to most RNA sequences prior to presentation of the RNA strand to the pore for sequencing. The current recommended protocol for DRS utilizes Induro^®^ RT and requires reverse transcription at 60°C. We demonstrate that these RT conditions promote hydrolysis of the RNA strand. We further show that UltraMarathonRT^®^ (uMRT), an ultraprocessive reverse transcriptase with intrinsic helicase activity that works optimally at 30°C, can be incorporated into a new uMRT-based DRS method that results in longer RNA reads in ONT DRS and longer final isoform predictions. We optimize this reaction along with other molecular biology steps and demonstrate the performance improvements of this new workflow on the benchmark sample, Universal Human Reference RNA, along with human brain RNA. This improved DRS protocol should empower new discoveries by the scientific community.

## Introduction

Oxford Nanopore direct RNA sequencing (DRS) profiles native RNA molecules rather than cDNA copies, thereby preserving endogenous base modifications and enabling strand-specific, isoform-resolved measurements in a single pass.^1,2^ Across a range of biological systems, DRS has supported transcriptome-wide isoform discovery and quantification, direct mapping of RNA chemical modifications, and measurement of 3′-end features such as poly(A)-tail length and cleavage sites.^1,3,4^ Since DRS sequencing initiates at the 3’ end of the RNA molecule, it is also less prone to internal priming artifacts seen in many cDNA sequencing methods.

These capabilities have advanced multiple areas of RNA biology. Viral and herpesvirus studies have resolved complex latency programs and promoter/TSS/TES architectures directly from native RNA.^5,6^ Native RNA sequencing has profiled protozoan and microbial transcriptomes as well as eukaryotic models interrogating the complexities of transcription and RNA processing.^7–9^ In human tissues, DRS has enabled isoform-level analyses of RNA modification landscapes in mRNAs and long noncoding RNAs.^10,11^ Specialized protocols further extend DRS to circular RNAs, miRNAs, and tRNAs.^12–14^

The higher error rate of DRS relative to long read cDNA sequencing along with the higher required input RNA mass have limited the widespread adoption of DRS technology. In addition, practical factors still limit routine DRS performance. Long and highly structured transcripts are often underrepresented due to RNA fragility and end truncation during library preparation, which depress read-length distributions and full-length coverage.^15^ Poly(A)-tail estimation is sensitive to processing and trimming parameters and therefore benefits from standardized workflows.^4^ Furthermore, modification calling remains chemistry- and context-dependent, requiring appropriate controls and calibration. Periodic updates to chemistry and basecalling models (e.g., RNA004 and associated Dorado models) necessitate iterative benchmarking to ensure cross-study comparability.^2,16^

In an effort to reduce the limitations of DRS, we reviewed the molecular biology workflow leading up to engagement with the nanopore and identified the synthesis of the RNA-cDNA hybrid by reverse transcriptase (RT) as a key step for improvement. We demonstrate that the RT buffer and temperature conditions currently recommended during DRS library preparation result in random hydrolysis of RNA. We develop an optimized workflow based on UltraMarathonRT^®^ (uMRT), an ultraprocessive, group II intron reverse transcriptase that possesses intrinsic helicase activity^17,18^ enabling it to work optimally at 30°C. We benchmark this workflow against the current ONT-recommended DRS workflow using two human samples and demonstrate increased representation of long transcripts. The resulting transcriptomes uncover more genes and isoforms with uMRT, improving the utility of DRS as a platform for discovery.

## Results

### uMRT Reaction Conditions Minimize RNA Degradation

In Oxford Nanopore direct RNA sequencing (ONT DRS), the RNA is first converted to an RNA-cDNA hybrid prior to presenting the RNA strand to the pore for sequencing. The current protocol recommends using the reverse transcriptase Induro^®^ for this conversion step.^19^ Induro requires incubation at 60°C, which we reasoned may increase the hydrolysis rate of the RNA due to the high temperature in the magnesium-containing RT buffer. To test hydrolysis rates of RNA during the RT reaction conditions, we incubated *in vitro* transcribed 7.6 kb RNA derived from the Hepatitis C viral genome (HCV) at the recommended temperature for uMRT, Induro and two commonly used mutants of MMLV RT, Superscript^™^ IV and Maxima^™^ (Figure 1A). After incubation of the RNA in 1X buffer for the recommended DRS RT time of 30 min, we found that 98% of the RNA was still intact in the uMRT conditions (30°C), while the other RT incubation conditions resulted in substantial RNA degradation ranging from ∼30-50%.

**Figure 1.**
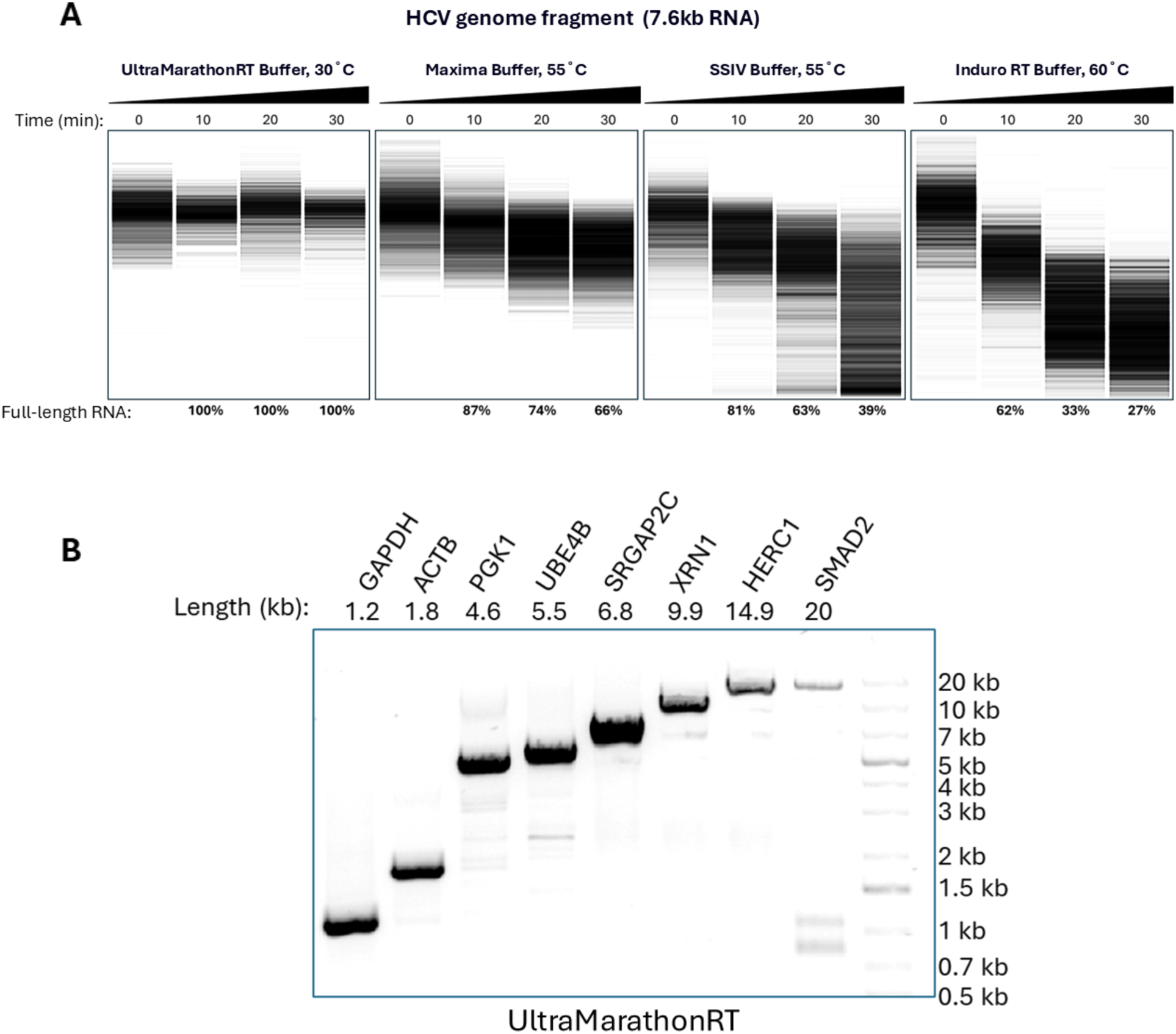
UltraMarathonRT’s low operating temperature preserves RNA integrity while synthesizing full-length cDNA. **(A)** Stability of a 7.6 kb HCV small genome RNA fragment was tested by incubation in the reaction buffers for UltraMarathonRT, Maxima H Minus, SuperScript IV, and Induro at their recommended reaction temperatures for 0, 10, 20, and 30 minutes, respectively. No reverse transcription was performed. The RNA samples were then analyzed using the Agilent™ 2100 Bioanalyzer™ system. **(B)** Amplification of human RNAs with lengths ranging from 1.2 kb to 20 kb using UltraMarathonRT Two-Step RT-PCR Kit. Oligo(dT) primed reverse transcription was performed using 100 ng of Hela total cellular RNA. PCR Amplification of cDNA was performed using gene-specific primers for GAPDH, ACTB, PGK1, UBE4B, SRGAP2C, XRN1, HERC1, & SMAD2.

Since the DRS RT step initiates cDNA synthesis at the polyA tail, we tested uMRT performance in generating long cDNAs by performing poly-dT primed cDNA synthesis at 30°C followed by PCR toward endogenous human genes (Figure 1B). The RT-PCR results showed detection of endogenous human genes ranging in size from 1.2 kb to 20 kb, consistent with uMRT’s well established high processivity. Given the low hydrolysis rate of RNA in uMRT reaction conditions and the inherent processivity of the enzyme, we hypothesized that uMRT may be an excellent RT for use in the DRS protocol.

### An Optimized Direct RNA-Seq Workflow with uMRT

In an effort to maximize RNA integrity and performance for DRS with uMRT as the engine of cDNA synthesis, we optimized several additional steps in the workflow leading up to Oxford Nanopore’s final proprietary adapter step that generates the ready-to-sequence library. Optimization of the library preparation protocol resulted in several deviations from legacy DRS methods including an RNA conditioning step to reduce intramolecular RNA interactions, a heat-cool annealing step for the RT adapter (RTA), changes in the ligase enzyme, bead purification between ligation and RT step, and addition of a quench buffer prior to the heat inactivation of the RT (Figure 2A).

**Figure 2.**
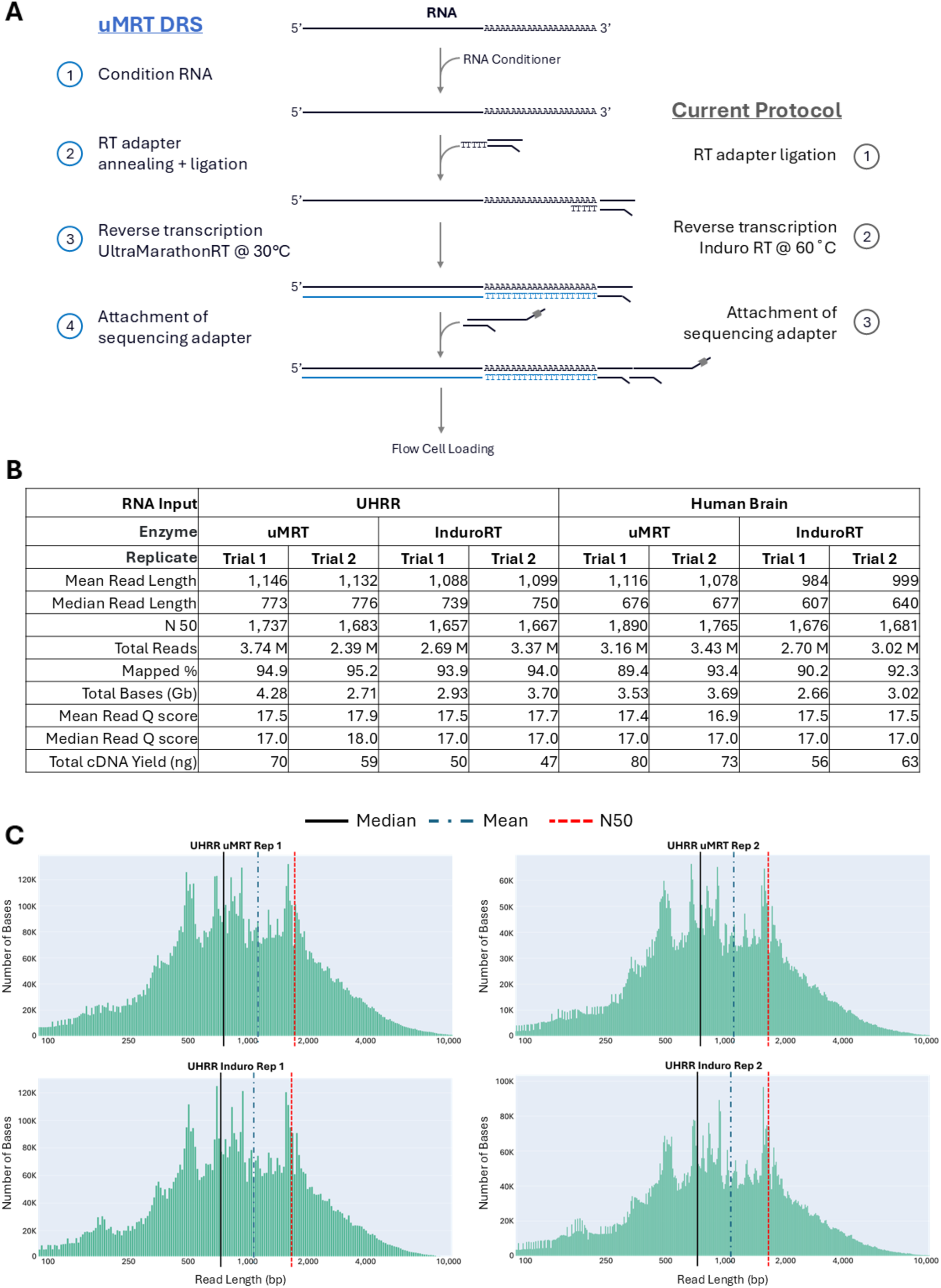
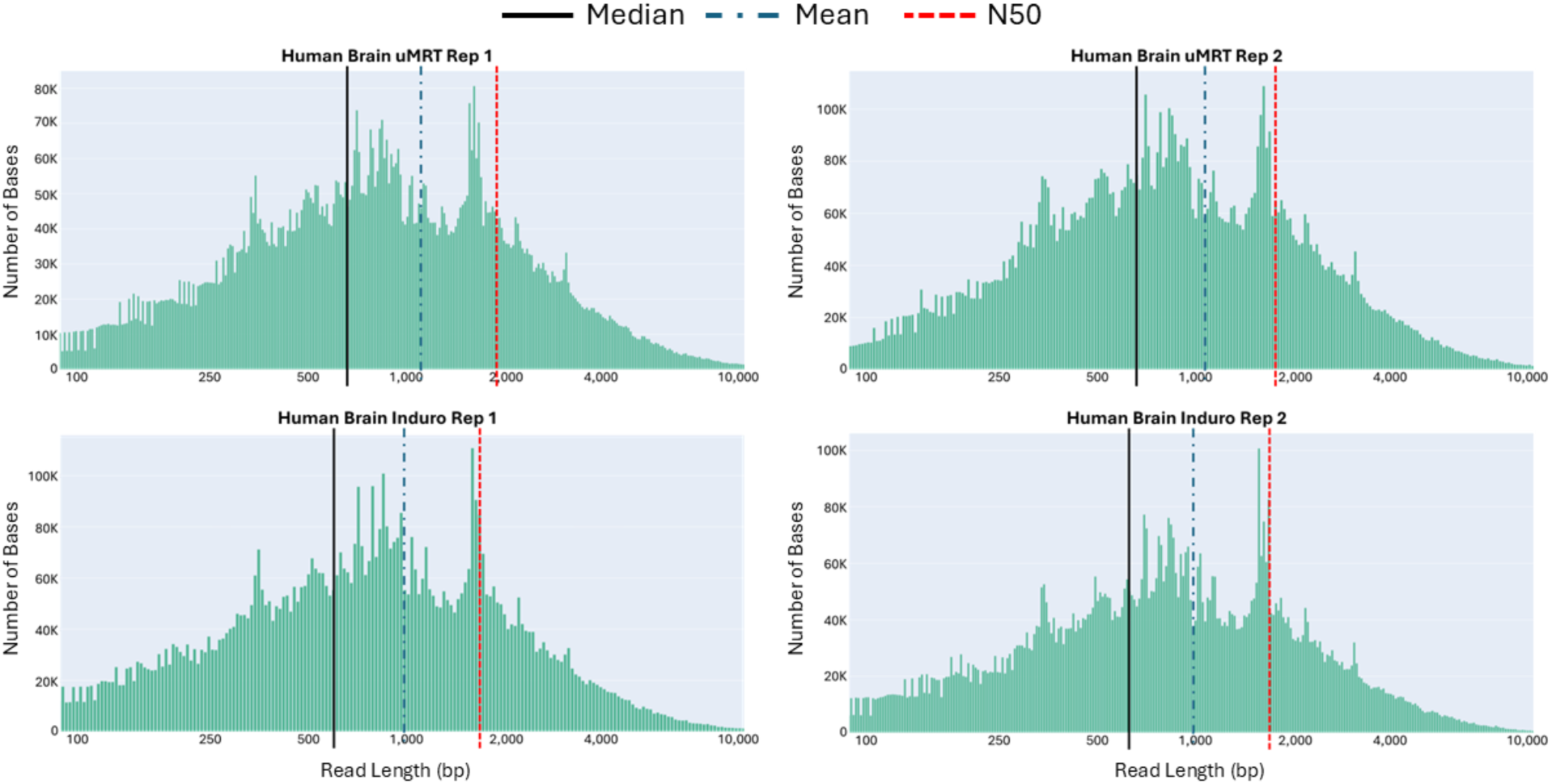
Direct RNA Sequencing (DRS) with UltraMarathonRT generates longer read lengths. **(A)** Overview of the workflow used to perform DRS with UltraMarathonRT compared to the current protocol. **(B)** Total read analysis and cDNA yield for DRS using 1 µg of total Universal Human Reference RNA (UHRR) and Human Brain RNA with uMRT and Induro. **(C)** Weighted log-transformed histograms of unmapped read lengths for UHRR and Human Brain DRS trials with uMRT and Induro.

To benchmark this optimized workflow versus the current recommended protocol from ONT, we chose two commercial total RNA sources: universal human reference RNA (UHRR) and human brain. UHRR is commonly used in large scale RNA-seq benchmarking studies both to test sample preparation methods and sequencing technologies.^20^ Human brain was chosen since it is rich in transcript complexity with alternative mRNA isoforms of a wide range in size.^21^

### Performance of an optimized uMRT-based workflow in Oxford Nanopore DRS

We prepared eight total libraries, including duplicates from both total RNA samples using the legacy Induro workflow and the optimized uMRT workflow. After the cDNA synthesis and cleanup steps, the samples were separately ligated to the ONT RNA004 adapter harboring the motor protein and quantified by fluorometry. The resulting RNA-cDNA hybrid yields were ∼25% higher with the optimized uMRT method (Figure 2B). Each library was then sequenced on individual ONT Minion flow cells.

The sequencing results revealed that mean read lengths for libraries prepared with uMRT showed modest improvements over the existing protocol including longer mean read length, median read length, and N50. Importantly, the new uMRT-based method did not impact the mapping rates or Q scores compared to the legacy methods (Figures 2B and S1A-C). The histograms for the RNA reads display similar overall topology with similar spikes in density, likely due to abundant RNAs that are present in each sample (Figures 2C,D).

In an effort to investigate the read-length differences and characterize isoform detection between the RT protocols, we combined the reads from each replicate and mapped the combined reads to the GRCh38 build of the human genome (Figure 3A). The total combined reads for libraries made with uMRT and Induro were nearly equivalent for UHRR (∼1% difference) so we proceeded in the analysis using all of the combined reads for each RT sample. For human brain, the difference in total combined reads were larger (∼13% more reads for the uMRT combined sample). Therefore, we randomly down sampled the total combined reads of uMRT to match the combined reads of Induro before mapping. The histograms for the mapped reads display similar topography to the histograms for the total reads in Figure 2C with an observed decrease in lengths under 100 bp for all samples. A slight shift in read length statistics was apparent in the uMRT conditions compared to Induro with the largest differences observed in the human brain. (Figure 3B upper panels versus lower).

**Figure 3.**
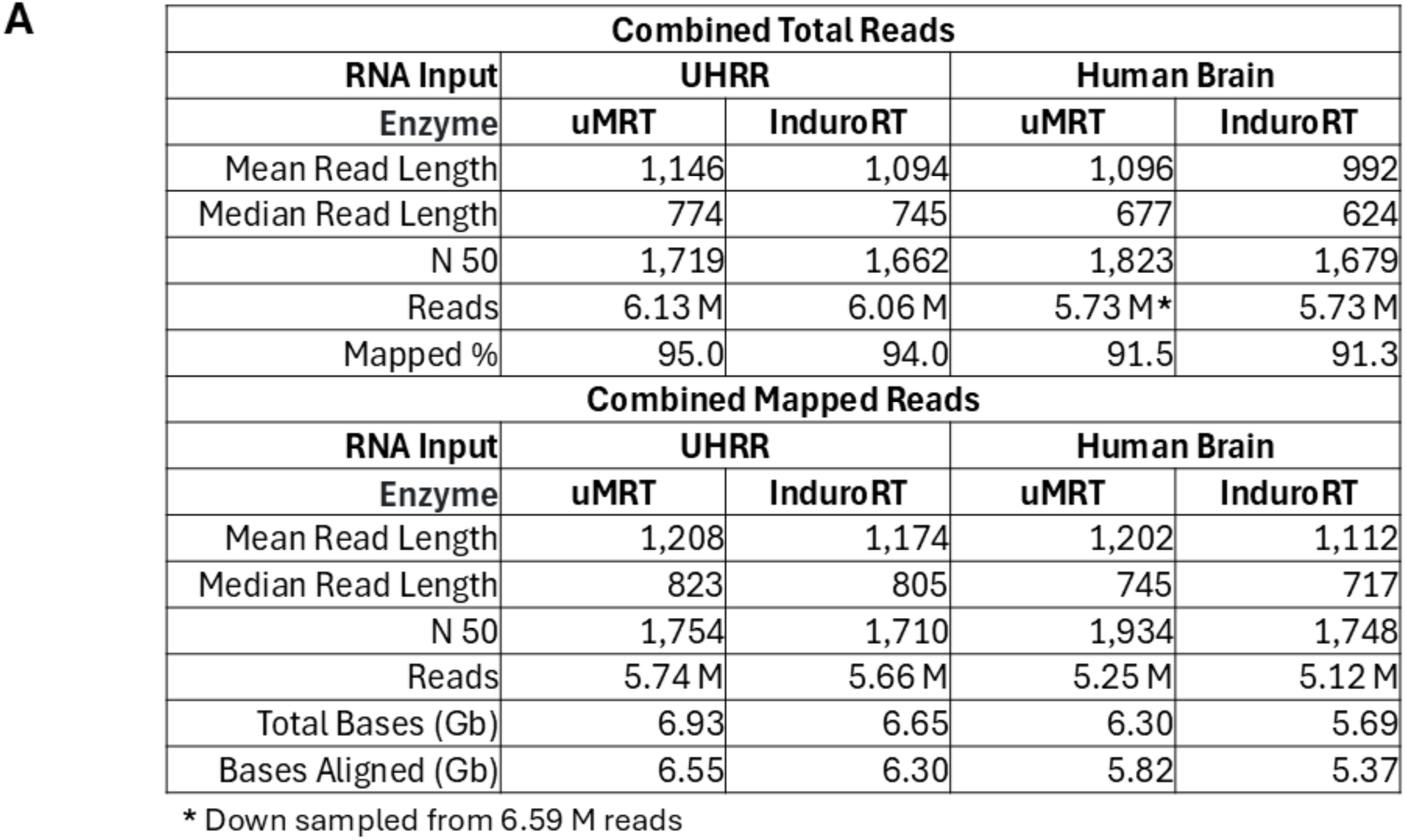
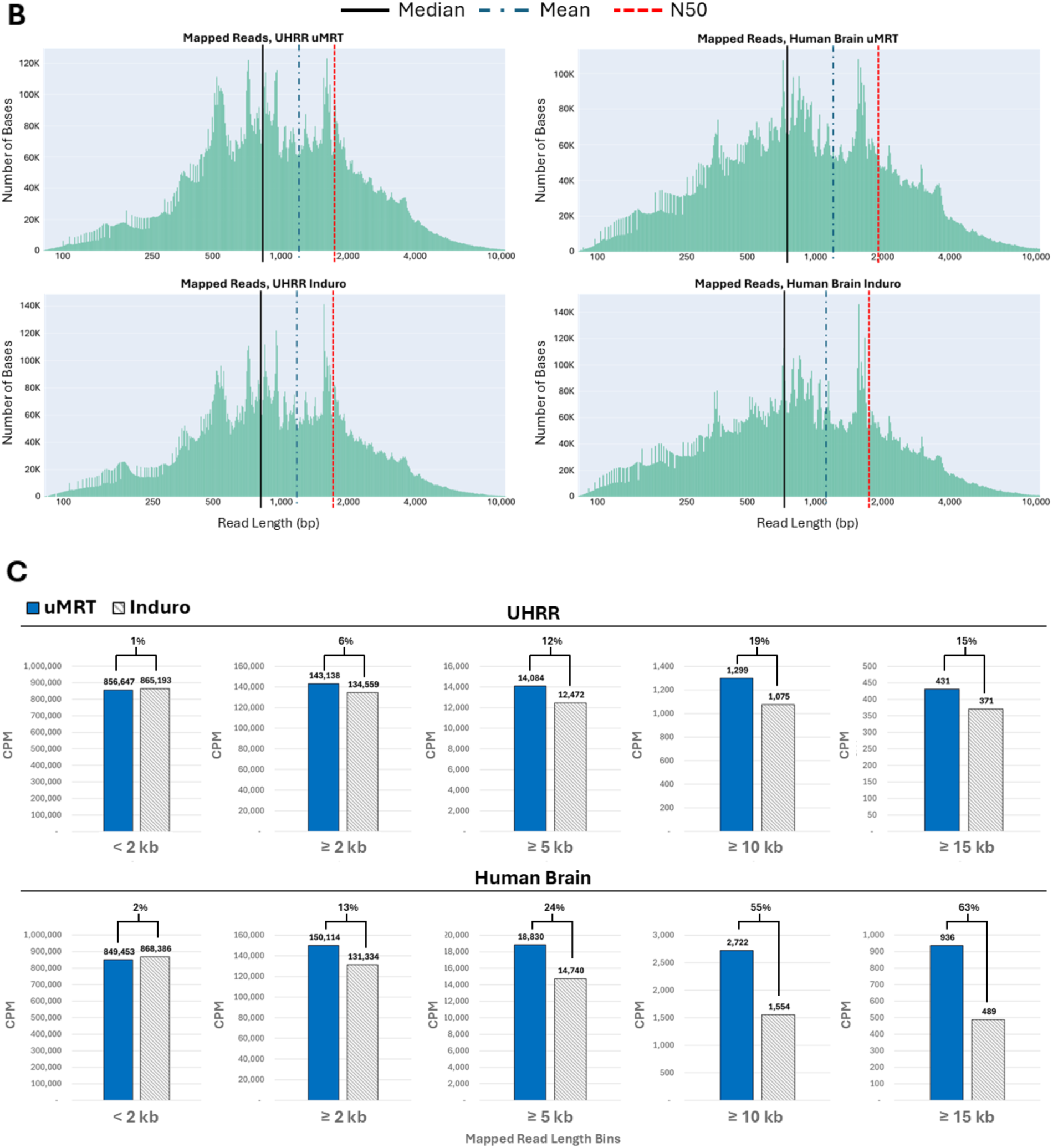
uMRT generates higher CPM’s of reads ≥ 2 kb. **(A)** Mapped read analysis of combined replicates for UHRR and Human Brain DRS with uMRT and Induro. Combined reads from Human Brain DRS samples for uMRT were down sampled from 6.59 million to 5.73 million reads using SEQTK. **(B)** Weighted log-transformed histograms of mapped read lengths for combined UHRR and Human Brain samples. **(C)** Mapped read Count Per Million (CPM) of combined UHRR and Human Brain samples. Percent difference between samples was calculated using the formula |x-y|/avg(x,y).

After observing this slight, but consistent overall advantage for the uMRT method, we segmented the total mapped reads into size bins to further investigate the effects of the new protocol on longer transcripts. Analysis of the binned RNA abundances as mapped read counts per million (CPM) uncovered a stepwise increase in counts as a function of transcript length (Figure 3C). While the protocols deliver similar CPM for smaller transcripts (<2 kb), performance improvements for the uMRT method increase progressively with longer read length bins. For instance, a modest shift is observed in the aggregate of all mapped read CPM ≥2 kb, but the difference becomes more dramatic when segmenting reads into ≥5 kb, ≥10 kb and ≥15 kb length bins. Interestingly, this segmentation uncovered a starker contrast between the two protocols for the human brain sample versus the UHRR, perhaps due to the higher prevalence of long mRNA isoforms in the human brain.

Since DRS is often used for isoform discovery and characterization, we compared the two workflows with respect to the number of detected genes and unique transcripts (also termed isoforms). For transcript analysis, we used two software packages; the ONT-recommended EPI2ME transcriptome workflow (<u>GitHub - epi2me-labs/wf-transcriptomes</u>) and IsoQuant^22^ which is optimized for long read RNA sequencing data from ONT and PacBio platforms.

In both methods, the resulting assembled transcripts were compared to the GENCODE v47 human annotations, an increasingly comprehensive catalog of transcript isoforms built from many human RNA samples and sequencing technologies.^23^

The EPI2ME software reported a slight advantage for the uMRT libraries in both detected genes and transcripts compared to Induro libraries for both the UHRR and human brain samples (Figure 4A). The analysis also showed that while the mean length of the transcripts from both libraries are relatively similar, the uMRT method provided greater detection of unique transcripts (Figure 4B). Agreeing with our earlier observations in mapped CPM, larger differences between the two methods were observed in longer transcript length bins for the human brain sample compared to the UHRR sample. For instance, the human brain transcriptome generated by the uMRT method contained nearly double the percentage of transcripts ≥15 kb (Figure 4C). When comparing the 10 longest human brain transcripts identified by both DRS methods, the 10^th^ longest transcript identified by uMRT was longer than longest transcript identified by Induro. When visually inspecting these using IGV, 7 of the 10 longest transcripts identified by the uMRT method contained a single continuous read of similar or nearly identical length to the corresponding transcript compared to only 3 using the Induro method (Figure 4D).

**Figure 4.**
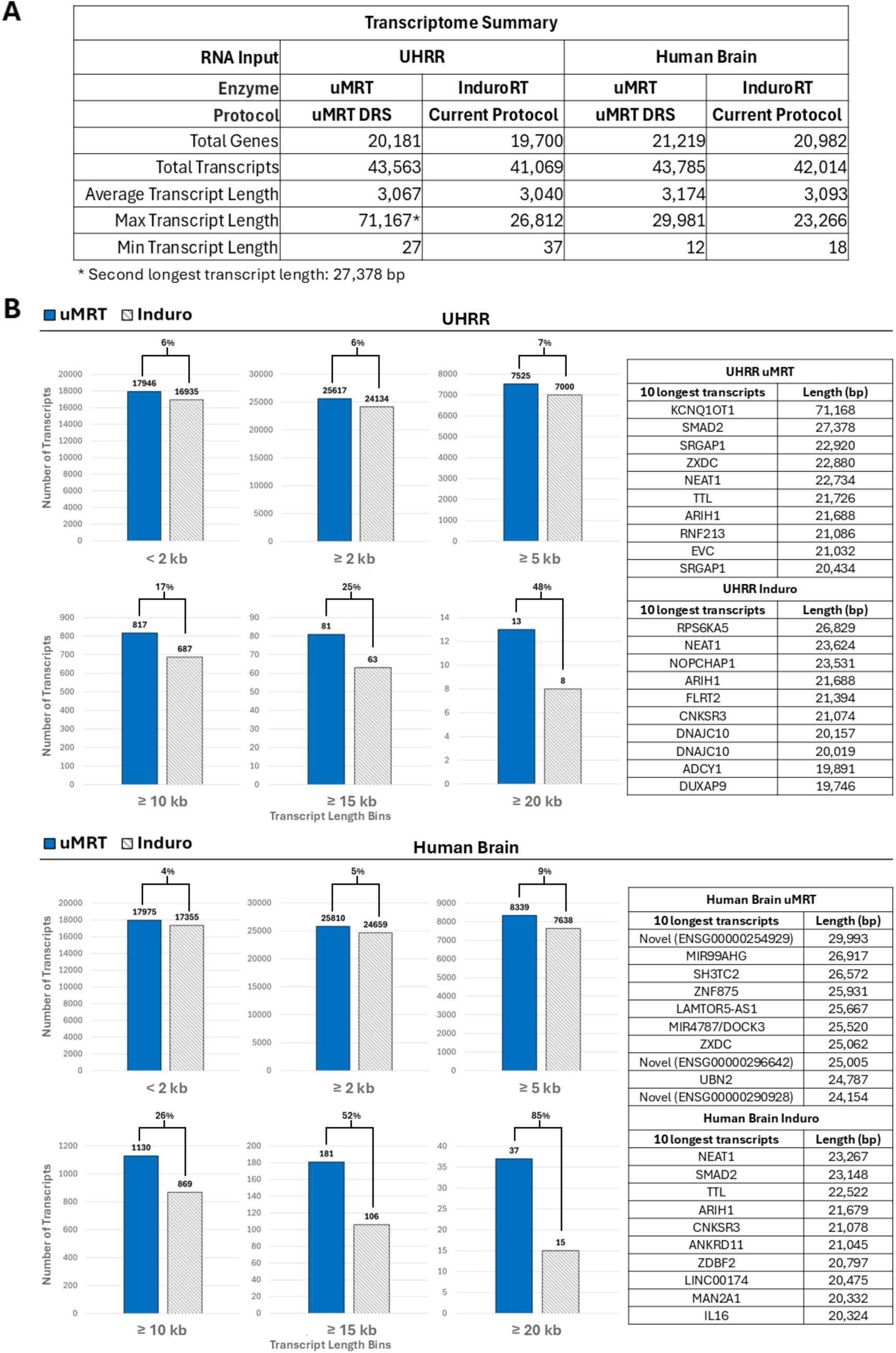
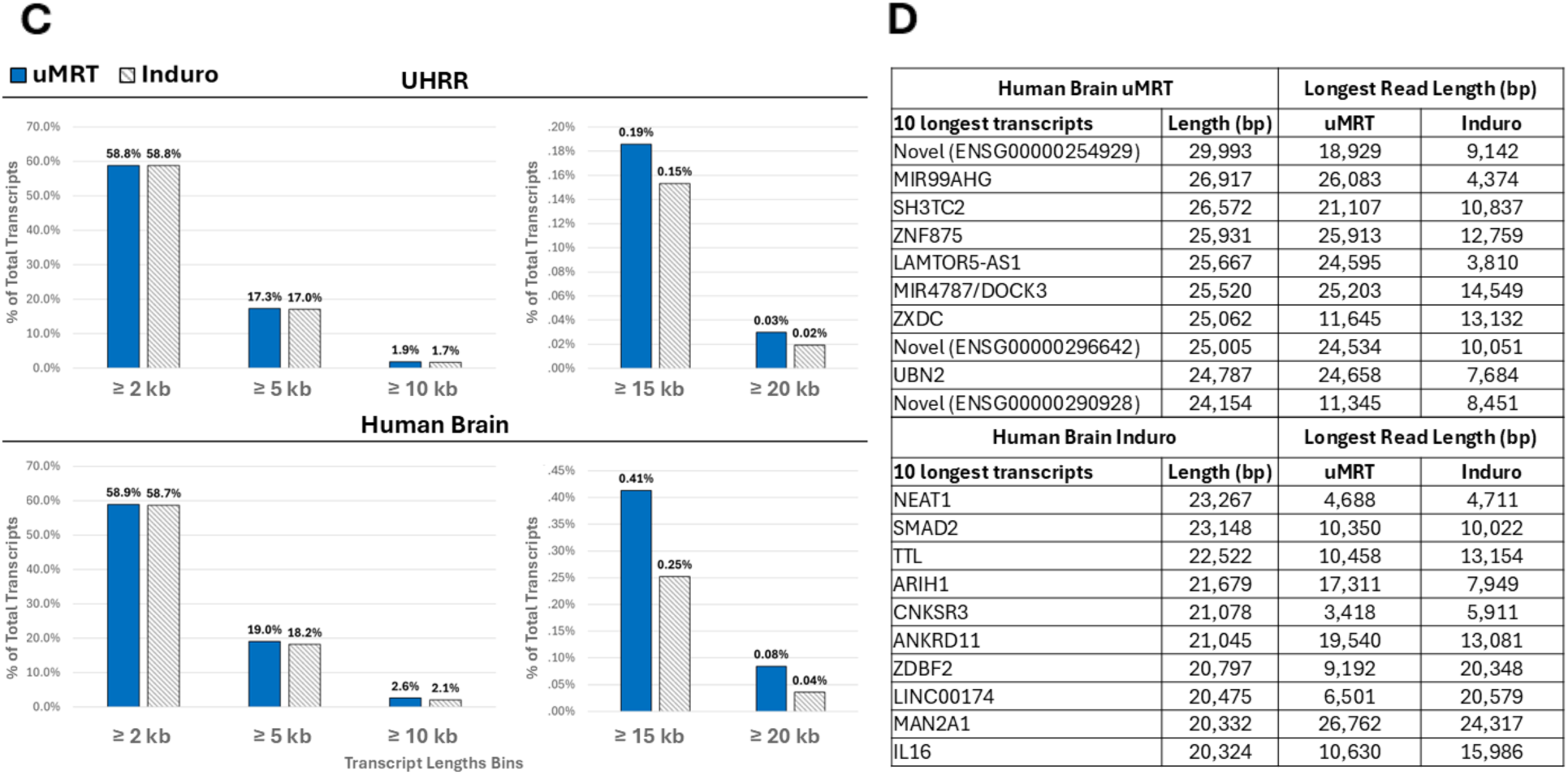
uMRT identifies more genes and longer transcripts. **(A)** Transcriptome analysis of the combined reads analyzed in Fig 3. Transcriptome was generated using the EPI2ME transcriptome workflow. **(B)** (left) Number of transcripts binned by transcript length. (right) 10 longest transcripts identified by uMRT and Induro. Parent gene of transcript is listed instead of specific transcript ID (see Supplementary Table 1 for all specific transcript ID’s). Percent difference between samples was calculated using the formula |x-y|/avg(x,y). **(C)** Percentage of total transcripts by varying bin lengths. **(D)** List of the longest continuous reads for each of the 10 longest transcripts identified in Human Brain samples. Longest reads were selected from the pool of same-strand reads aligned to the parent gene of the target transcript using IGV.

### Characterization and classification of Isoforms

To gain deeper insight into isoform characterization outside the capabilities of the EPI2ME software, we also generated transcript models using IsoQuant due to its reported performance on DRS data in terms of precision and recall. Benchmarked against other long read analysis tools, IsoQuant reports the highest percentage of transcripts confirmed by at least three other tools, while also reporting the lowest false discovery rate amongst the tools.^22^ The resulting IsoQuant annotations for the uMRT and Induro libraries indeed reflected this higher stringency with slightly lower numbers of total genes and transcripts detected versus EPI2ME (Figure 5A). However, the pattern remains consistent wherein more total genes and transcripts are seen in uMRT libraries as compared to libraries made with Induro. We then classified the IsoQuant transcripts using SQANTI3, comparing the transcripts to existing isoform annotations from GENCODE. This tool is especially powerful for annotating the chains of splicing events along a predicted transcript model and characterizing whether these events are novel. ^24^

**Figure 5.**
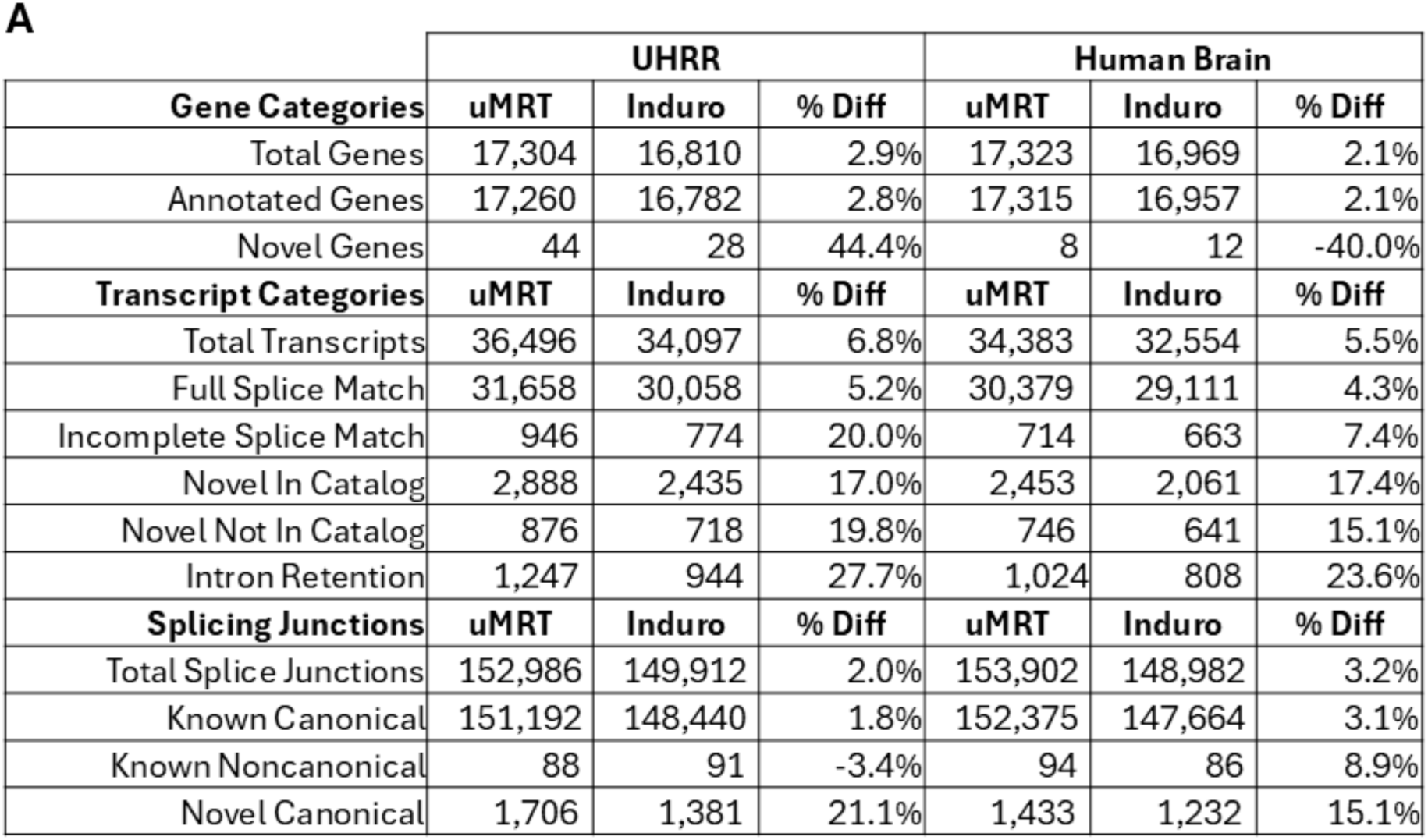
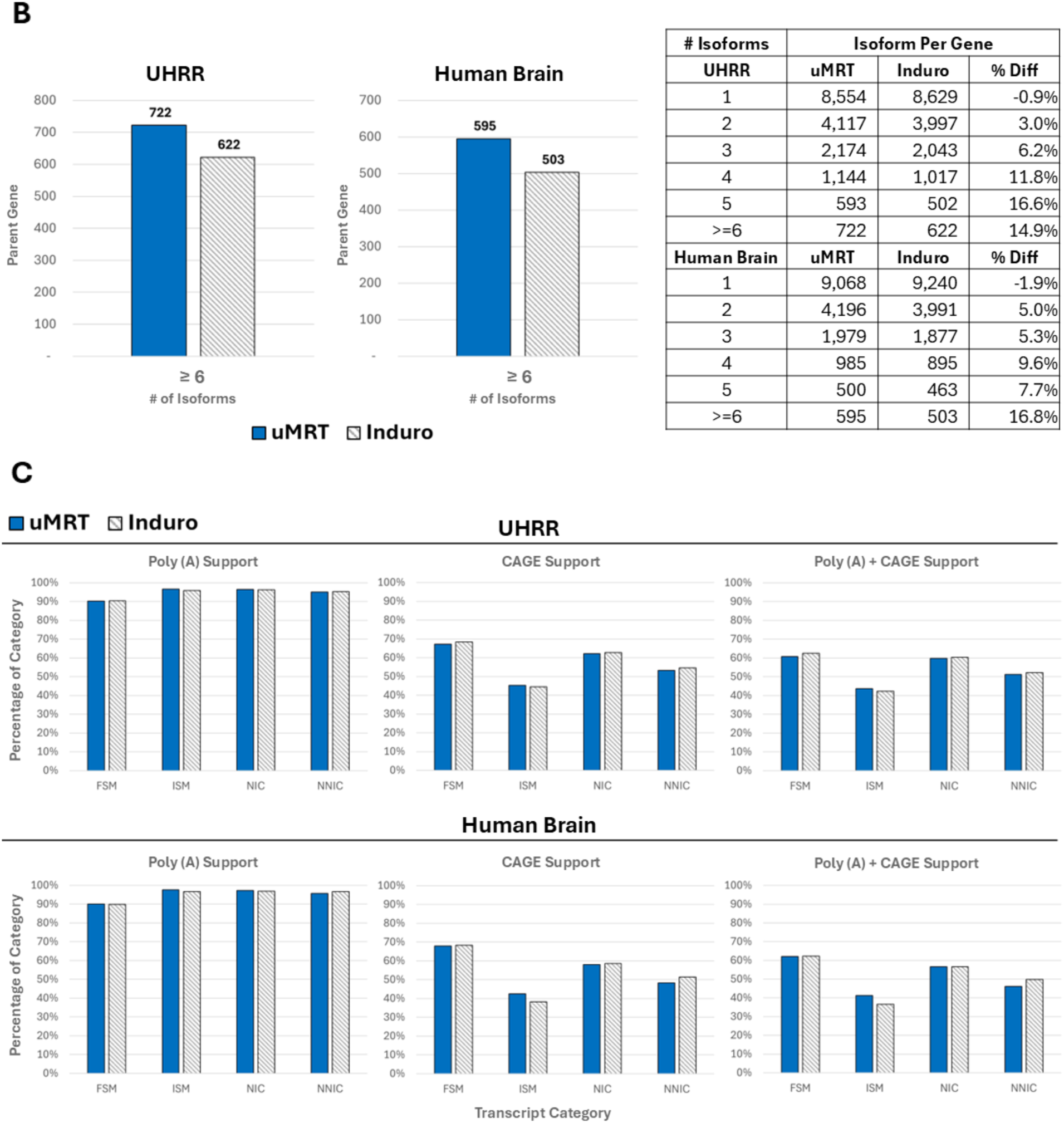
uMRT identifies more novel isoforms and intron retention. **(A)** Comparison of genes and transcripts identified by uMRT and Induro generated by IsoQuant and SQANTI3. **(B)** Comparison of isoforms per gene identified by uMRT and Induro generated by SQANTI3. Percent difference (% Diff) was calculated using the formula |x-y|/avg(x,y). Negative values for % Diff represent instances where Induro generated a higher value than uMRT. **(C)** Comparison of 5’ CAGE, PolyA, and 5’CAGE + PolyA Support for different transcript categories in UHRR and Human Brain generated by SQANTI3. Full Splice Match (FSM), Incomplete Splice Match (ISM), Novel In Catalog (NIC), Novel Not In Catalog (NNIC).

SQANTI3 further enabled us to characterize both canonical and non-canonical splicing events while leveraging CAGE data marking 5’ transcript start sites (TSS) as well as polyA motifs marking 3’ cleavage/polyadenylation sites.

The SQANTI3 results uncovered modest advantages in the Full Splice Match (FSM) and Incomplete Splice Match (ISM) isoforms for uMRT libraries with both RNA samples (Figure 5A). Of particular interest in long read RNA sequencing analyses are the novel isoform categories, since these can encode new proteoforms. The uMRT libraries from both RNA samples identified ∼17% more Novel in Catalog (NIC) and 15-20% more Novel Not in Catalog (NNIC) isoforms than Induro libraries. The NIC events represent known splice sites but used in a new pattern (e.g. skipping of an exon previously thought to be constitutive), while NNIC events represent completely novel splice sites not previously seen in a reference annotation. In comparing and classifying the total splice junctions, uMRT libraries detected 15-20% more novel canonical splice sites. Additionally, the number of transcripts detected with an intact intron (intron retention) was 24-28% higher in the uMRT libraries, consistent with the longer transcript lengths for these molecules since they still contain one or more unspliced retained or detained introns. Finally, we found that the uMRT libraries had more isoforms per gene for both the UHRR sample and the human brain sample (Figure 5B).

Consistent with prior DRS studies, the gene body coverage for both UHRR and human brain displayed 3’ bias and was nearly identical between uMRT and Induro (Supplementary Figure S1D). This may indicate that RNA integrity can degrade over the course of the 24-72 hour DRS sequencing run. Comparison of the detected isoforms with orthogonal data types indicate that the polyA motif is present in the vast majority of the isoforms (>90%) for both uMRT and Induro libraries, consistent with the sequencing mechanism starting at polyA tails (Figure 5C). Short read CAGE data, which marks 5’ ends of transcripts, varies more between the different categories of SQANTI isoform classification, and is largely matched between the uMRT and Induro methods as well.

## Discussion

Direct RNA sequencing (DRS) delivers unique access to native RNA to characterize modifications and view the RNA processing landscape at the full-length isoform level. Many transcripts in plants and animals are especially complex and can be 10’s of kilobases in length, challenging even the most robust sequencing technologies. We hypothesized that the fragile nature of RNA would make the library preparation leading up to the pore engagement especially important. Incubation of RNA in a magnesium-containing buffer at pH >8 accelerates RNA hydrolysis, yet these are the conditions that are essential for all reverse transcriptases as they copy RNA. We show that the current RT conditions used for DRS with Induro can degrade RNA due to the high temperature (60°C) recommended for the reaction. This led us to test uMRT, a reverse transcriptase that can unwind and copy long RNA at 30°C where hydrolysis is significantly reduced.

In addition, we carried out extensive optimization of other steps leading up to the proprietary ONT motor protein ligation. The optimized protocol introduces (i) an RNA conditioning step to reduce intramolecular RNA interactions, (ii) a heat-cool-anneal step to better deliver the RT adapter (RTA) to the polyA tail, (iii) a ligase swap, (iv) a bead purification between ligation and RT to remove inhibitory carryover, and (v) a quench buffer before RT heat inactivation. In all, these changes increase the library yield. Interestingly, the optimized uMRT workflow consistently produced more than double the ONT recommended minimum library yield for sequencing (30 ng). Because of this, we were recently able to successfully perform DRS using only 500ng of total RNA, and 100 ng of poly(A)+ RNA as starting input amounts (Supplementary Figure S2E). Though the potential for using half the RNA input addresses an underlying limitation of DRS, more work needs to be done to confirm its reproducibility.

Protecting RNA from hydrolysis during library preparation appears to generate a modest but reproducible shift toward longer reads. Across two human RNA samples (UHRR and brain), libraries prepared with the uMRT workflow showed higher mean and median read lengths and a longer N50, while maintaining mapping rates and per-read quality (Q-scores). The distribution shifting toward longer reads with uMRT is consistent with stabilization of more long isoform molecules during library construction.

Binning analyses by transcript length demonstrated that both protocols perform similarly for shorter RNAs (<2 kb), while the uMRT protocol led to a progressive enrichment of counts in larger size bins, with the gains most evident at >10 kb and >15 kb, where DRS has historically under-represented transcripts. The effect is more pronounced in human brain than in UHRR, likely due to the known prevalence of long isoforms in cell types found in the brain.

We show that the length-distribution improvements translate into incremental but meaningful gains in discovery power. Using EPI2ME, uMRT libraries detected more genes and more isoforms than the Induro workflow in both samples, with brain again showing the largest benefit. IsoQuant, a stricter, benchmark-validated caller, confirmed the same pattern despite reporting fewer total isoform models. Comparison of the isoforms to annotations with SQANTI3 showed modest boosts in the number of isoforms across classification categories. Notable gains were observed in novel in-catalog (NIC) and novel not-in-catalog (NNIC) events, more novel canonical splice junctions, and a marked increase in intron-retention calls in the uMRT libraries. More isoforms per gene were also observed in the less stringent EPI2ME transcriptome analysis (Supplementary Figure S2A). Collectively, this additional information augments the potential mRNAs and, in turn, the encoded proteoforms translated in these samples.

We expect this optimized DRS library preparation approach to benefit any new projects using DRS since the workflow is simple and the reagents for this preparation are now commercially available in kit form. The data obtained from this optimized DRS approach should be compatible and comparable to prior data obtained using the SQK-RNA004 chemistry, but with the benefit of more reads at the longer end of the transcript spectrum.

Long read RNA sequencing continues to provide new insights into biology and the complexities of transcription and post-transcriptional process. We foresee that this optimized method will be especially useful for groups studying long isoforms or genes exhibiting high splicing complexity and will result in new biological insights. Neurobiology, stem cell biology, or developmental biology, along with cancer transcriptomics, will benefit from high quality full-length isoform predictions and novel splice junction discoveries to help unlock new biology. Long read cDNA sequencing on PacBio and ONT platforms may also benefit from the stabilization effect seen here, along with the excellent processivity with uMRT.

Future work will benefit from sequencing additional tissues and input amounts, along with testing uMRT in adaptations of DRS geared toward non-polyadenylated RNA (e.g. tRNA, miRNA). Extending the evaluation to chemistries and callers not assessed here, and incorporating orthogonal controls where appropriate, will help calibrate the discovery gains with biological truth. We anticipate that the stabilization of RNA will always aid in these pursuits, but the gain seen here via library preparation does not erase the inherently fragile nature of RNA as a biopolymer. RNA preparation methods and sequencing methods such as the 24-72 hour nanopore runs used in this study can contribute to RNA degradation as well. Here, we show that changing the RT step to a lower temperature helps preserve long RNA, but safeguarding the RNA from inside the cell through to the final sequencing output file is a goal to continue to strive forward with future innovations.

## Methods and Materials

### RNA Stability Assay

An *in vitro* transcribed 7.6 kb HCV small genome RNA fragment was incubated in 1X reaction buffer for UltraMarathonRT^®^ (RNAConnect^™^ Cat. No. R1002), Maxima^™^ H Minus (ThermoScientific^™^, Cat#K1652), SuperScript^™^ IV (Invitrogen^™^, Cat#18091300), and Induro^®^RT (NEB^™^, Cat#M0681) at their recommended reaction temperatures for 0, 10, 20, and 30 min respectively. For Induro RT, incubation temperature of 60°C was used following the ONT Direct RNA sequencing protocol. Samples were then analyzed using the Agilent^™^ RNA 6000 Pico Kit (PN# 5067-1513) on a 2100 Bioanalyzer^™^ following manufacturers protocol.

Band intensities were obtained using Image J (v1.54p) and binned according to intervals of RNA ladder sizes ∼0.2 – 3.9 kb (degraded RNA) and ∼4.0 – 10.0 kb (full-length RNA). Percentages were calculated by dividing the binned band intensity by the total band intensity. RNA stability was determined by changes in binned percentages over time. (N=2)

### RT-PCR targeting full-length human genes of varying lengths

RT-PCR was performed on 100 ng of HeLa total cellular RNA (Takara^™^, Cat#636543) using UltraMarathonRT^®^ Two-Step RT-PCR Kit (RNAConnect^™^, Cat#R1005) according to the manufacturer recommendations. Reverse transcription was primed using oligo(dT)_18_ to copy all mRNA. PCR Amplification of cDNA was performed using gene-specific primers for GAPDH, ACTB, PGK1, UBE4B, SRGAP2C, XRN1, HERC1, & SMAD2 (Supplementary Table 3). PCR product was subjected to gel electrophoresis using a 0.8% agarose gel, then visualized using GelDoc Go imaging system (Bio-Rad).

### Oxford Nanopore Direct RNA Sequencing

Direct RNA sequencing was performed on 1 μg of Total Universal Human Reference RNA (Agilent^™^, Cat#750500) and Total Human Brain RNA (Invitrogen^™^, Cat#AM7962) using the Oxford Nanopore Technologies (ONT) Direct RNA Sequencing kit (ONT, Cat#SQK-RNA004).

Libraries were generated using two different protocols. Both protocols required the ONT SQK-RNA004 kit to perform direct RNA sequencing as it contains the oligo adapters, motor protein, and flow cell loading reagents.

#### Library preparation using the current protocol

The complete library preparation protocol currently recommended by Oxford Nanopore Technologies (ONT) can be found here:

https://nanoporetech.com/document/direct-rna-sequencing-sqk-rna004#overview-of-the-protocol

In short, the ONT reverse transcription adapter is first ligated to poly(A) tailed RNA using NEB^™^ T4 DNA Ligase (NEB^™^, cat#M0202M). Reverse transcription is then performed using Induro^®^RT (NEB^™^, Cat#M0681) at 60°C for 30 minutes followed by 70°C for 10 minutes to inactivate enzyme. The cDNA/RNA duplex is then purified using RNAClean XP beads (Beckman Coulter^®^, Cat#A63987). A second ONT oligo adapter bound to the ONT motor protein is then ligated to the first RNA-ligated adapter followed by a final bead purification.

#### Library preparation using UltraMarathonRT

Direct RNA Sequencing libraries for uMRT were generated using the UltraMarathonRT Direct RNA Sequencing kit (RNAConnect^™^, Cat#R1008) following the manufacturer recommended protocol:

https://cdn.shopify.com/s/files/1/0872/5613/8010/files/UltraMarathonRT_Direct_RNA_Sequencing_Protocol_11.18.25.pdf?v=1765295646

In short, the total RNA is first conditioned using an RNA conditioning reagent. The ONT reverse transcription adapter is then annealed to poly(A) tailed RNA at 55°C for 1 minute followed by a snap chill to 4°C. The annealed adapter is then ligated to the RNA using a DNA Ligase. The ligated RNA is then purified using RNAClean XP beads (Beckman Coulter^®^, Cat#A63987). Reverse transcription is performed using UltraMarathonRT at 30°C for 30 minutes. A quenching buffer is added to the reaction before heat inactivation at 70°C for 5 minutes followed by bead purification of the cDNA/RNA duplex. A second ONT oligo adapter bound to the ONT motor protein is then ligated to the first RNA-ligated adapter followed by a final bead purification.

For uMRT only, DRS was also performed using 500 ng of the Total Human Brain RNA previously mentioned, as well as 300 ng and 100 ng of Human Skeletal Muscle Poly A+ RNA (Takara^™^, Cat#636120).

Library yields were measured using the Qubit 1X dsDNA HS Assay Kit (ThermoScientific^™^, Cat#Q33231) on a Qubit 4 Fluorometer (ThermoScientific^™^).

### MinION flow cell loading and sequencing

The DRS library yields were loaded on a FLO-MIN004RA flow cell and sequenced using a MinION device for 48-72 hours, or until ∼3-3.5 million reads were obtained.

### Direct RNA Sequencing Data Analysis

#### Basecalling

Raw POD5 files were base called and converted to FASTQ using Dorado (v.7.8.3) via the ONT MinKNOW software with RNA004 – 4kHz chemistry, super-accurate basecalling type (≥ Q10), and v5.1.0 – 130bps base calling model.

#### Mapping and read length analysis

Total reads for each individual sample were aligned to the human genome GRCh38.p14 using the EPI2ME alignment workflow (v1.2.4) and minimap2^25^ (v2.28). FASTQ files for replicate UHRR and Human Brain sequencing runs were also combined using OmicsBox^26^ (v3.5). For uMRT Human Brain sample, total combined reads were down sampled from 6.59 to 5.73 million reads using SEQTK (v1.3-r106). Read length and quality for all samples were analyzed using EPI2ME alignment workflow (v1.2.4), Nanoplot^27^ (v1.46.1) and LongQC^28^ (v1.2). Mapped read count per million (CPM) was calculated using the number of binned read lengths generated by NanoPlot. Mapped gene body coverage plots and read quality by length plots for combined UHRR and Human Brain samples were generated using RSeQC^29,30^ (v5.0.1).

#### Transcriptome and isoform analysis

Transcriptomes were created using the EPI2ME transcriptome workflow (v1.7.1) and IsoQuant^22^ (v3.6.2). In brief, the combined FASTQ files for replicate DRS runs were mapped to the human genome GRCh38.p14 and aligned using the GENCODE v47 transcript annotations using minimap2^25^ (v2.28). The transcript models generated by IsoQuant were further characterized using SQANTI3^24,31^ (v3.5.2) using additional reference TSS annotation and PolyA motif files supplied in the GitHub repository (see “SQANTI3” in the “Code Availability” section).

Inspecting the longest reads for the 10 longest transcripts in Human Brain samples was performed using IGV (v2.19.6). First, the locus of the parent gene of the target transcript was visualized using IGV. Next, reads were grouped by strand and sorted by aligned read length. The longest continuous read within the pool of same-strand aligned reads was selected as the longest read. The BAM and index files generated via the EPI2ME transcriptome workflow for uMRT and Induro were used in this IGV analysis. The 10 longest transcripts for each respective DRS workflow were generated by the EPI2ME transcriptome workflow.

## Data Availability

Sequencing data generated for this study are available at the NCBI Sequence Read Archive through the BioProject accession number PRJNA1378338.

## Code Availability

EPI2ME Alignment Workflow: https://github.com/epi2me-labs/wf-alignment

EPI2ME Transcriptome Workflow: https://github.com/epi2me-labs/wf-transcriptomes

SQANTI3: https://github.com/ConesaLab/SQANTI3

OmicsBox: https://www.biobam.com

## Supplementary Tables

Supplementary Table 1: Complete transcript table generated by EPI2ME

Supplementary Table 2: Complete transcript table generated by SQANTI3

Supplementary Table 3: Primers used in RT-PCR

## Acknowledgments

This work was supported in part by grants from the National Institutes of Health (R01HG011868 to BRG and R44GM153078 to RNAConnect).

