## Supplementary Figures for "Improved long-transcript representation in Oxford Nanopore direct RNA sequencing with UltraMarathonRT"

**Figure S1.**

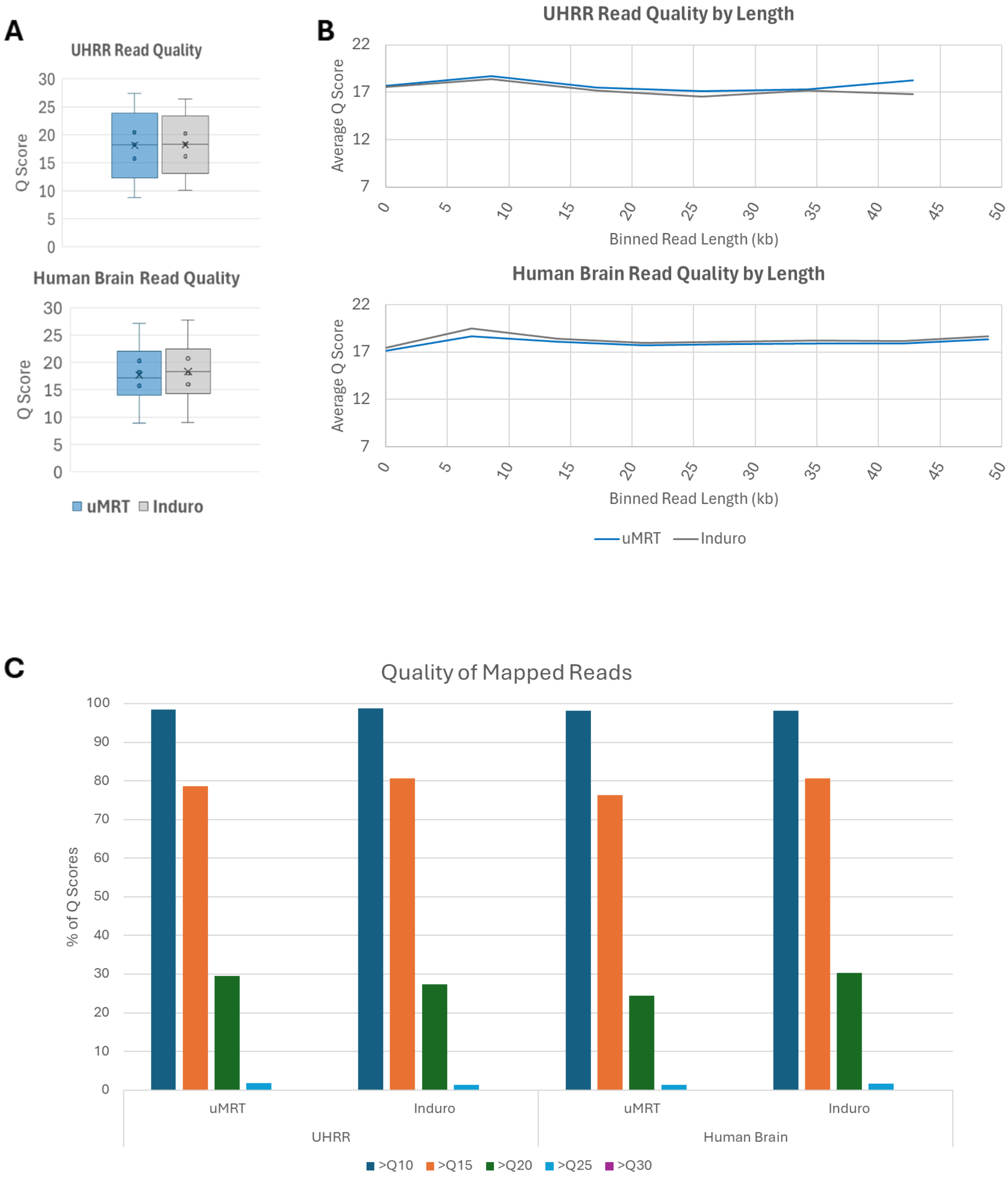
A
UHRR Read Quality
Q Score
30
25
20
15
10
5
0
Human Brain Read Quality
Q Score
uMRT Induro
B
UHRR Read Quality by Length
Average Q Score
22
17
12
7
0
5
10
15
20
25
30
35
40
45
50
Binned Read Length (kb)
Human Brain Read Quality by Length
Average Q Score
Binned Read Length (kb)
uMRT Induro
C
Quality of Mapped Reads
% of Q Scores
100
90
80
70
60
50
40
30
20
10
0
uMRT
Induro
uMRT
Induro
UHRR
Human Brain
>Q10 >Q15 >Q20 >Q25 >Q30

**D**

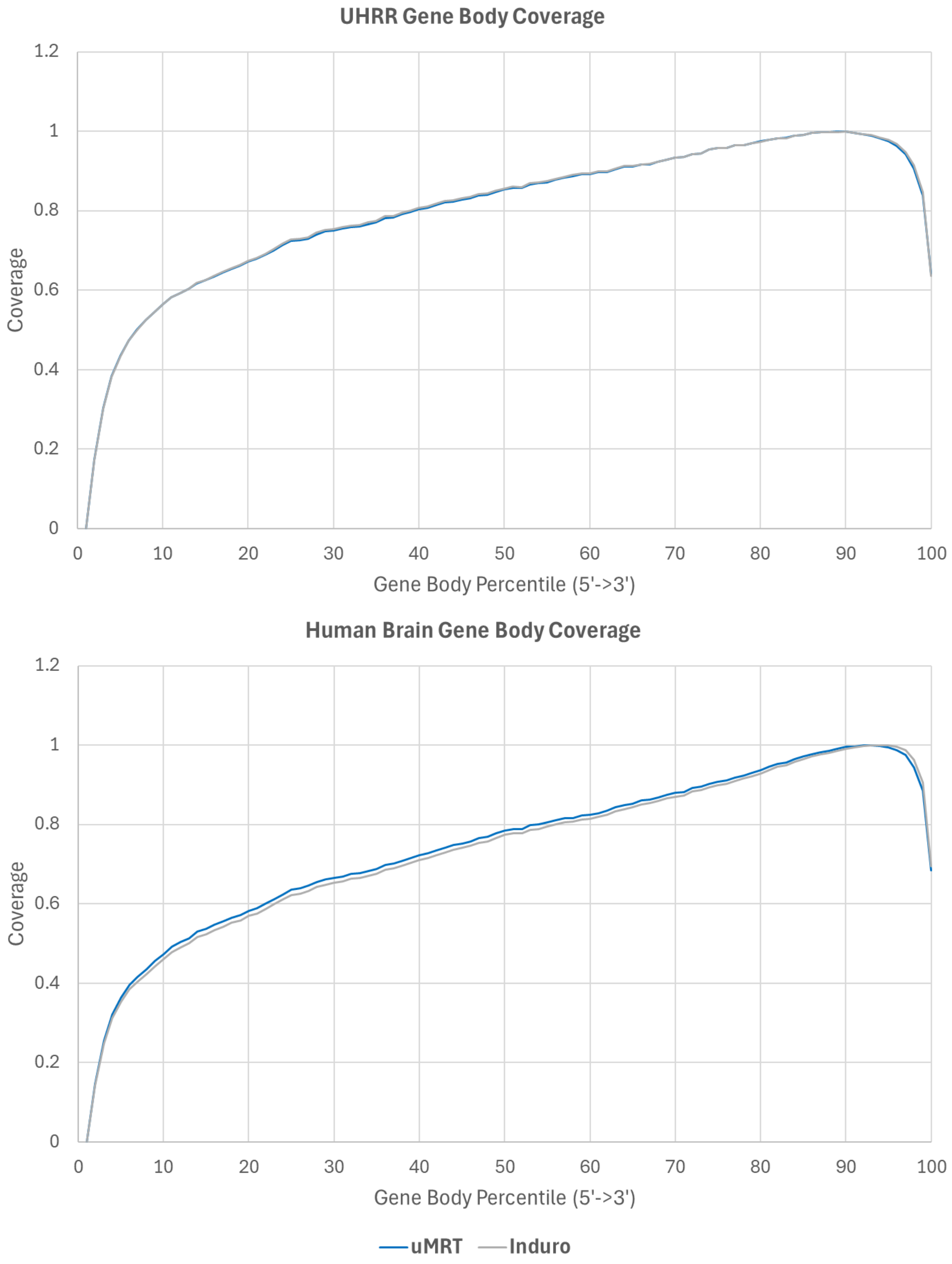


**Figure S1. Quality and gene body coverage of combined mapped reads. (A)** Box-whisker plot of Q scores and **(B)** plot of average Q score by length for total reads in UHRR (top) and Human Brain (bottom) generated by LongQC. **(C)** Percentage of Q scores for mapped reads in UHRR (left) and Human Brain (right) generated by Nanoplot. **(D)** Gene body coverage of both enzymes in UHRR and Human Brain RNA.

**Figure S2.**

**A**

| # of Isoforms | Isoforms Per Gene | | |
|---|---|---|---|
| **UHRR** | **uMRT** | **Induro** | **% Diff** |
| Unkown | 81 | 52 | 43.6% |
| 1 | 9747 | 9706 | 0.4% |
| 2 | 4426 | 4453 | -0.6% |
| 3 | 2456 | 2362 | 3.9% |
| 4 | 1396 | 1296 | 7.4% |
| 5 | 766 | 697 | 9.4% |
| 6 | 454 | 392 | 14.7% |
| 7 | 263 | 210 | 22.4% |
| 8 | 166 | 127 | 26.6% |
| 9 | 96 | 80 | 18.2% |
| ≥ 10 | 118 | 97 | 19.5% |

| # of Isoforms | Isoforms Per Gene | | |
|---|---|---|---|
| **Human Brain** | **uMRT** | **Induro** | **% Diff** |
| Unkown | 257 | 226 | 12.8% |
| 1 | 10125 | 10285 | -1.6% |
| 2 | 4544 | 4523 | 0.5% |
| 3 | 2434 | 2321 | 4.8% |
| 4 | 1348 | 1297 | 3.9% |
| 5 | 722 | 720 | 0.3% |
| 6 | 454 | 386 | 16.2% |
| 7 | 264 | 196 | 29.6% |
| 8 | 153 | 125 | 20.1% |
| 9 | 79 | 76 | 3.9% |
| ≥ 10 | 132 | 118 | 11.2% |

**B**

| | UHRR | | Human Brain | |
|---|---|---|---|---|
| | **uMRT** | **Induro** | **uMRT** | **Induro** |
| Total Assignments | 5,983,264 | 5,832,648 | 5,175,248 | 5,105,791 |
| Assignment Poly-A Percentage | 50.5 | 42.3 | 49.0 | 42.4 |
| Known Transcript Models | 31,782 | 30,163 | 30,472 | 29,159 |
| NIC Transcript Models | 3,484 | 2,928 | 2,876 | 2,477 |
| NNIC Transcript Models | 4,197 | 3,633 | 4,296 | 3,879 |
| Ambiguous Reads | 1,496,343 | 1,437,767 | 1,202,470 | 1,152,491 |
| Inconsistent Reads | 235,238 | 198,097 | 195,046 | 166,536 |
| Intergenic Reads | 8,102 | 7,572 | 20,127 | 18,695 |
| Noninformative Reads | 117,915 | 172,929 | 148,211 | 160,207 |
| Unique Reads | 2,882,009 | 2,898,828 | 2,492,785 | 2,539,174 |
| Unique with minor difference Reads | 314,094 | 296,297 | 227,027 | 226,003 |

**C**

| | UHRR | | | Human Brain | | |
|---|---|---|---|---|---|---|
| **Transcript Categories** | **uMRT** | **Induro** | **% Diff** | **uMRT** | **Induro** | **% Diff** |
| FSM | 31,658 | 30,058 | 5.2% | 30,379 | 29,111 | 4.3% |
| ISM | 946 | 774 | 20.0% | 714 | 663 | 7.4% |
| NIC | 2,888 | 2,435 | 17.0% | 2,453 | 2,061 | 17.4% |
| NNIC | 876 | 718 | 19.8% | 746 | 641 | 15.1% |
| Genic Genomic | 14 | 11 | 24.0% | 22 | 9 | 83.9% |
| Antisense | 36 | 17 | 71.7% | 6 | 10 | -50.0% |
| Fusion | 62 | 70 | -12.1% | 61 | 57 | 6.8% |
| Intergenic | 16 | 14 | 13.3% | 2 | 2 | 0.0% |
| Genic Intron | - | - | | - | - | |
| Total Transcripts | 36,496 | 34,097 | 6.8% | 34,383 | 32,554 | 5.5% |
| | **UHRR** | | | **Human Brain** | | |
| **Isoform SubCategories** | **uMRT** | **Induro** | **% Diff** | **uMRT** | **Induro** | **% Diff** |
| Internal Fragment | 8 | 11 | -31.6% | 7 | 8 | -13.3% |
| 3' Fragment | 336 | 267 | 22.9% | 270 | 291 | -7.5% |
| 5' Fragment | 346 | 290 | 17.6% | 229 | 198 | 14.5% |
| At Least 1 novel splice site | 802 | 670 | 17.9% | 670 | 583 | 13.9% |
| Annotated Splice Junction | 1,428 | 1,306 | 8.9% | 1,259 | 1,067 | 16.5% |
| Annotated Splice Site | 567 | 463 | 20.2% | 478 | 432 | 10.1% |
| Mono Exon | 1,611 | 1,447 | 10.7% | 1,660 | 1,499 | 10.2% |
| Intron Retention | 1,247 | 944 | 27.7% | 1,024 | 808 | 23.6% |
| Multi Exon | 112 | 96 | 15.4% | 75 | 64 | 15.8% |
| Ref Match | 30,049 | 28,613 | 4.9% | 28,721 | 27,614 | 3.9% |

**D**

| | Total | | | |
|---|---|---|---|---|
| | **UHRR** | | **Human Brain** | |
| **Cage Support** | **uMRT** | **Induro** | **uMRT** | **Induro** |
| 5'Cage Support | 24,006 | 22,862 | 22,768 | 21,709 |
| PolyA Support | 33,182 | 31,037 | 31,290 | 29,624 |
| CAGE + PolyA Support | 21,992 | 20,958 | 20,921 | 19,979 |
| Total Transcripts | 36,496 | 34,097 | 34,383 | 32,554 |
| | **UHRR** | | **Human Brain** | |
| **Cage Support %** | **uMRT** | **Induro** | **uMRT** | **Induro** |
| 5'Cage Support | 65.8% | 67.0% | 66.2% | 66.7% |
| PolyA Support | 90.9% | 91.0% | 91.0% | 91.0% |
| CAGE + PolyA Support | 60.3% | 61.5% | 60.8% | 61.4% |

**E**

| Read Summary | | | |
|---|---|---|---|
| **RNA Input** | **Poly(A)+ Skeletal Muscle** | | **Total Human Brain** |
| **Input Level** | **300 ng** | **100 ng** | **500 ng** |
| Mean Read Length | 962 | 953 | 915 |
| Median Read Length | 718 | 745 | 5,587 |
| N 50 | 1,430 | 1,410 | 1,600 |
| Total Reads | 2.66 M | 3.42 M | 2.06 M |
| Mapped % | 92.2 | 94.6 | 90.7 |
| Aligned Reads | 2.45 M | 3.24 M | 1.87 M |
| Total Aligned bases (Gb) | 2.56 | 3.26 | 1.88 |
| Mean Read Quality (Q score) | 16.9 | 17.7 | 16.8 |
| Median Read Quality (Q score) | 17.0 | 18.0 | 17.0 |
| Total cDNA Yield (ng) | 55 | 33 | 34 |
| **Transcriptome Summary** | | | |
| **RNA Input** | **Poly(A)+ Skeletal Muscle** | | **Total Human Brain** |
| **Input Level** | **300 ng** | **100 ng** | **500 ng** |
| Total Genes | 12,226 | 13,237 | 16,741 |
| Total Transcripts | 18,869 | 21,503 | 28,110 |
| Average Transcript Length | 2,503 | 2,625 | 2,777 |
| Max Transcript Length | 21,153 | 21,142 | 23,172 |
| Min Transcript Length | 31.0 | 20.0 | 13.0 |

**Figure S2. Additional transcript metrics (A)** Isoform per gene generated by EPI2ME transcriptome workflow. Percent difference (% Diff) was calculated using the formula |x-y|/avg(x,y). Negative values for % difference represent instances where Induro generated a higher value than uMRT. **(B)** Transcript metrics generated by IsoQuant. **(C)** Complete transcript metrics generated by SQANTI3. Percent difference (% Diff) was calculated using the formula |x-y|/avg(x,y). Negative values for % difference represent instances where Induro generated a higher value than uMRT. **(D)** CAGE and PolyA support for total transcripts generated by SQANTI3. **(E)** Alignment and transcriptome analysis for DRS with uMRT using 300 ng and 100 ng Skeletal Muscle Poly(A)+ RNA and 500 ng of total Human Brain RNA generated by EPI2ME.
